# Bulk RNA sequencing deconvolution of pancreatic ductal adenocarcinoma identifies cancer-associated fibroblast subsets associated with survival and tumor microenvironment composition

**DOI:** 10.64898/2026.04.03.716260

**Authors:** Nicole Dam, Mischa F.B. Steketee, Gaby Strijk, Willem de Koning, Lukas J.A.C. Hawinkels, Vera Kemp, Casper H.J. van Eijck, Yongsoo Kim, Casper W.F. van Eijck, Bram W. van Os

**Affiliations:** Department of Cell and Chemical Biology, Leiden University Medical Centre, Leiden, The Netherlands; Department of Gastroenterology-Hepatology, Leiden University Medical Centre, Leiden, The Netherlands; Department of Pathology, Cancer Center Amsterdam, Amsterdam UMC, location VUmc, Amsterdam, The Netherlands; Department of Surgery, Erasmus MC Cancer Institute, University Medical Centre Rotterdam, Rotterdam, The Netherlands; Department of Pulmonary Medicine, Division of Solid Tumor Immunology Research Rotterdam (STIRR), Erasmus MC Cancer Institute, University Medical Centre Rotterdam, Rotterdam, The Netherlands; Department of Pathology and Clinical Bioinformatics, Erasmus MC Cancer Institute, University Medical Centre Rotterdam, Rotterdam, The Netherlands

## Abstract

Pancreatic ductal adenocarcinoma (PDAC) is a highly lethal cancer characterized by a high abundance of cancer-associated fibroblasts (CAFs), which influence therapy response, tumor biology and tumor aggressiveness. CAFs are a heterogeneous cell type and previous single-cell RNA sequencing (scRNAseq) of PDAC tumors identified three main CAF subtypes: myofibroblastic, inflammatory and antigen-presenting CAFs (myCAF, iCAF, apCAF, respectively). However, scRNAseq on large patient cohorts is often not feasible due to costs and technical constraints. Therefore, bulk RNAseq deconvolution can be used to identify cell types within the heterogeneous tumor microenvironment. Here, Statescope deconvolution was used to identify different cell types of the tumor microenvironment within an early onset PDAC cohort, comprising 74 patients aged under 60. Three CAF populations were identified (iCAFs, myCAFs and desmoplastic CAFs), and their correlations with tumor microenvironment components, mutational signatures and survival were examined. iCAFs were associated with classical-like tumor cells, whereas myCAFs and desmoplastic CAFs correlated with basal-like tumor cells. Desmoplastic CAFs are associated with inflammatory granulocytes/neutrophils, while negatively associating with monocyte-derived macrophages and immature/transitional B cells. No associations were observed between mutational signatures and the abundance of CAF and epithelial tumor subtypes. Interestingly, a high abundance of CAFs, and specifically increased iCAF abundance, was associated with improved survival. This iCAF-mediated survival effect was predominantly apparent in female patients. All in all, deconvolution of bulk RNA sequencing data, followed by its integration with clinical and biological parameters, reveals the heterogeneity and prognostic implications of CAF subpopulations in the tumor microenvironment of early onset PDAC patients.

## Introduction

Pancreatic ductal adenocarcinoma (PDAC) is a highly aggressive cancer with a 5-year overall survival (OS) rate of only 13% in all stages combined [1]. Early symptoms of the disease are often absent, which complicates diagnosis of the disease. Consequently, approximately 90% of PDAC cases are identified at locally advanced or metastatic stages, making treatment more difficult. [2]. Patients often receive chemotherapy, typically combining 5-fluorouracil, leucovorin, irinotecan and oxaliplatin (FOLFIRINOX) or gemcitabine with nab-paclitaxel, sometimes alongside surgery [3, 4]. However, response rates to chemotherapy are low, with only around 17% to 47% of metastatic patients responding to FOLFIRINOX in different trials [5, 6]. Immunotherapies have shown limited benefit in treating PDAC, due to its immunologically cold tumor microenvironment (TME), which features a low abundance of anti-tumor directed immune cells and a high abundance of immunosuppressive cells [7].

A major reason for the limited success of current chemo- and immune directed therapies in PDAC, which also contributes to its immune-cold environment, is the pronounced stromal content. Up to 80% of PDAC tumor mass consists of non-tumor cells, comprising immune cells, endothelial cells, nerves, extracellular matrix (ECM) and cancer-associated fibroblasts (CAFs). Among these, CAFs are considered the most prominent component, due to their central role in mediating tumor growth, immune responses and treatment resistance [8]. Furthermore, desmoplasia, the excessive deposition of ECM, is a common feature of PDAC and directly results from the high density of CAFs [9]. This dense ECM network reduces vascularization and limits immune cell infiltration into the tumor. While CAF depletion was proposed as a strategy to counteract this, subsequent studies have shown that removing all CAFs can adversely affect patient survival [10]. This has led to the recognition that CAFs are a heterogeneous cell population, with both tumor-promoting and tumor-suppressing roles. Efforts to classify CAF subtypes have identified three well-defined and acknowledged subtypes of CAFs: myofibroblastic CAFs (myCAFs), inflammatory CAFs (iCAFs) and antigen-presenting CAFs (apCAFs) [11, 12]. myCAFs are distinguished by high expression levels of α-smooth muscle actin (α-SMA) and play a key role in ECM production and remodeling [11]. iCAFs secrete inflammatory cytokines, such as interleukin (IL)-6, and shape immune signaling within the stroma [11]. apCAFs are defined by their expression of MHC class II molecules and can therefore present antigens on their cell-surface [12]. Importantly, each subtype exhibits both anti-tumor and pro-tumor functions, suggesting further heterogeneity within these CAF subsets.

Typically, CAF subtyping has relied on the use of single-cell RNA sequencing (scRNAseq). However, its widespread application is constrained by high costs, limited sequencing depth and bias towards certain cell types introduced by the tissue dissociation and cell isolation protocols. Consequently, identifying CAF subsets has often been restricted to small datasets. While bulk RNA sequencing avoids many of these limitations, it lacks single-cell resolution. Deconvolution bridges this gap by integrating these methods. Deconvolution models are trained on scRNAseq datasets and applied to bulk RNAseq data to infer cell-type composition and cell type-specific gene expression profiles. This enables the separation of CAF subsets, as well as subsets of other cell types.

In this study, we used Statescope to deconvolute bulk RNAseq data, which allows estimation of cell-type abundance and their transcriptional subsets, referred to as cell states [13, 14]. Deconvolution was performed on bulk RNA sequencing data from a recently published cohort of 74 PDAC patients under 60 years of age, including 50 resected tumors from untreated patients and 24 resected tumors that received neoadjuvant FOLFIRINOX [15]. Statescope deconvolution of PDAC tumors previously identified five CAF subsets, alongside other cell types present in the TME, thereby facilitating analysis of intratumor heterogeneity [13, 14]. The current study differs from the previous study by using a cohort of early-onset PDAC cancers, which comprise around 15% of the patients in the Netherlands and which more frequently undergo resection and have better OS compared to patients above 60 years [16]. Furthermore, treatment regimens of the patients in this cohort were more uniform compared to the cohort used for the previously performed Statescope study. The aim of this project was to identify CAF subsets present within the PDAC tumors of this early onset patient cohort and to explore their associations with other cell types in the TME, the mutational signature of the tumors and clinical survival data.

## Material and methods

### Patient cohort

Data from whole exome and whole transcriptome sequencing of primary PDAC tumors from 74 patients under 60 years old were collected in a previously published study [15]. Inclusion criteria included resectable primary PDAC tumors with available DNA and RNA sequencing data. Exclusion criteria encompassed patients with high tumor mutational burden (TMB-H) or microsatellite instability (MSI), those who received neoadjuvant radiation or chemotherapy other than FOLFIRINOX, and patients with pancreatic tumors other than PDAC. Sequencing was performed on formalin-fixed, paraffin-embedded (FFPE) tissue, which was selected for tumor purity according to the following requirements: ≥10% tumor content for WTS or ≥20% for WES, and minimally 50mm² of tumor tissue, as described previously [15].

### Deconvolution

#### Alignment of DNA sequencing data and estimating tumor purity

Matched DNA fastq files containing pair-ended reads were aligned to the human reference GRCh38.p14 genome using BWA-MEM, streamed to samtools and sorted into coordinate-sorted BAM files. The reference genome was downloaded from the University of California, Santa Cruz (UCSC) genome browser. Aligned reads were saved in sorted BAM files and PCR/optical duplicates were marked with Picard (3.4.0).

Tumor purity was estimated from CNVkit (0.9.12) copy number profiles. Segmented CNVkit files were parsed to autosomal chromosomes and small segments were removed. A Gaussian mixture model (scikit-learn, 1.7.1) was used to identify peaks associated with losses and gains. Purity was inferred from how far the loss/gain log2 peaks are shifted away from the (assumed) diploid baseline and clonal single-copy losses (1 copy) and gains (3 copies) in the tumor.

#### Alignment of RNA sequencing data

Paired-end RNA-seq FASTQs were aligned to human GRCh38 using STAR (v2.7.11b) in two-pass mode with the GENCODEv46, GRCh38 reference FASTA with matching GTF annotation. The resulting raw count table was used as input for Statescope.

#### Statescope deconvolution

RNA raw count matrices and DNA-based tumor purities estimates were imported into Python, and duplicate gene entries were removed by summing across them to yield one row per gene symbol. Statescope (1.0.6) was initialized with the PDAC signature for 7 cell types: fibroblasts, epithelial cells, endothelial cells, NK(T) cells, mast cells, myeloid cells and B/plasma cells. The epithelial fraction was clipped at 0.99. Deconvolution was performed with GPU acceleration (CUDA 12.9) and expression refinement was executed twice, followed by State Discovery [13, 14]. The Statescope results were loaded into R Studio (2025.05.1, R version 4.5.1). Figures other than heatmaps were plotted with ggplot2 (4.0.0), heatmaps were created with either pheatmap (1.0.13) and Complexheatmap (2.24.1). Immune, endothelial and cancer-associated fibroblast (CAF) subsets were annotated by matching the genes with the highest positive and negative state loadings to known subsets in literature. Epithelial subsets were defined as classical or basal-like (or a mixture of the two) based on correlation-based scoring with the gene set described by Hilmi *et al*. 2025 [17]. To functionally annotate the epithelial cells, gene set enrichment analysis was performed using fgsea (1.34.2), with minSize = 5, and maxSize = 500, eps = 1^-10^, input gene ranked on stateloading, and gene sets downloaded from the msigdbr (25.1.1). Top regulated pathways were used to functionally annotate the epithelial subsets. Subset proportions were first transformed using a centered log-ratio (CLR) transformation. Each CLR-transformed subset was then regressed on tumor purity, and Spearman correlations were calculated on the resulting residuals.

### Statistical analyses

All analyses, except Kaplan-Meier curves and baseline characteristics, were performed in R Studio. Kaplan-Meier curves and baseline characteristics were analyzed using Graphpad Prism (10.2.3, San Diego, CA, USA). Baseline characteristics were checked for normal distribution (Shapiro-Wilk test) and statistical tests were adjusted accordingly. For numerical baseline characteristics data, Mann-Whitney U tests were performed, while categorical data were compared using Fisher’s exact test. All other statistical tests are defined in the figure legends, including multiple testing analyses. P values lower than 0,05 were considered significant.

## Results

### Patient cohort and identification of cell types using deconvolution of bulk RNAseq

A total of 74 PDAC patients with available DNA and bulk RNA sequencing data were retrieved from a cohort comprising early-onset PDAC cancer patients under 60 years old [15]. Baseline characteristics indicate that the median age at diagnosis was 55 years and 58% of the patients were male. Approximately a third of patients were treated with neoadjuvant FOLFIRINOX (**Table 1**). To identify the different cell types present in the TME, bulk transcriptomics data were deconvoluted using Statescope, followed by state discovery of these subsets. Tumor purity estimates derived from copy number analysis of DNA-seq data were used as prior information on the epithelial compartment during deconvolution. To assess the quality of the deconvolution, the fraction of the epithelial transcriptional signal identified through Statescope was compared to the estimated tumor purity through CNVkit, which showed a strong correlation (R^2^=0,91) and a linear relationship, supporting the validity of the deconvolution. The deviation from the identity relation (x = y) suggests that these tumor cells contributed disproportionally more to the transcriptional bulk than the other cell types. (**Supplementary figure 1A**).

**Table 1.** Baseline characteristics of included patients, n (%)

| Total patients = 74 |  |
| --- | --- |
| Sex, male | 43 (58%) |
| Age diagnosis (median, IQR) | 55 (50 - 57) |
| CA 19-9 at diagnosis (median, IQR) | 74,5 (28,75-244,5) |
| Differentiation grade |  |
| *Well* | 7 (9%) |
| *Moderate* | 29 (39%) |
| *Poor* | 20 (27%) |
| *Not assessed* | 18 (24%) |
| Neoadjuvant chemotherapy, FOLFIRINOX | 24 (32%) |
| Adjuvant chemotherapy |  |
| *FOLFIRINOX* | 54 (73%) |
| *Gemcitabine* | 3 (4%) |
| *Gemcitabine + Capecitabine* | 1 (1%) |
| *None* | 16 (22%) |
| Disease stage at end of follow-up |  |
| *(borderline) resectable* | 0 (0%) |
| *Locally advanced* | 0 (0%) |
| *Metastatic* | 25 (34%) |
| *Resected* | 45 (61%) |
| *Local recurrence* | 4 (5%) |
| Death | 39 (53%) |

*IQR: interquartile range*
*NGS: next-generation sequencing*

The cell types identified in the patient samples using Statescope include mast cells, epithelial cells, T and NK cells, B and plasma cells, myeloid cells, endothelial cells and cancer-associated fibroblasts (CAFs) (**Figure 1A**). Transcriptionally, epithelial cells (comprising primarily tumor cells) were most abundant, as expected given the tumor-purity selection criteria for the FFPE tissues used for sequencing. Although transcriptional signals of mast, T and NK cells were detected, they comprised a small fraction. Additionally, the abundance of B and plasma cells, myeloid cells, endothelial cells and CAFs varied strongly across patients (**Figure 1A, Supplementary table 1**).

**Figure 1.**
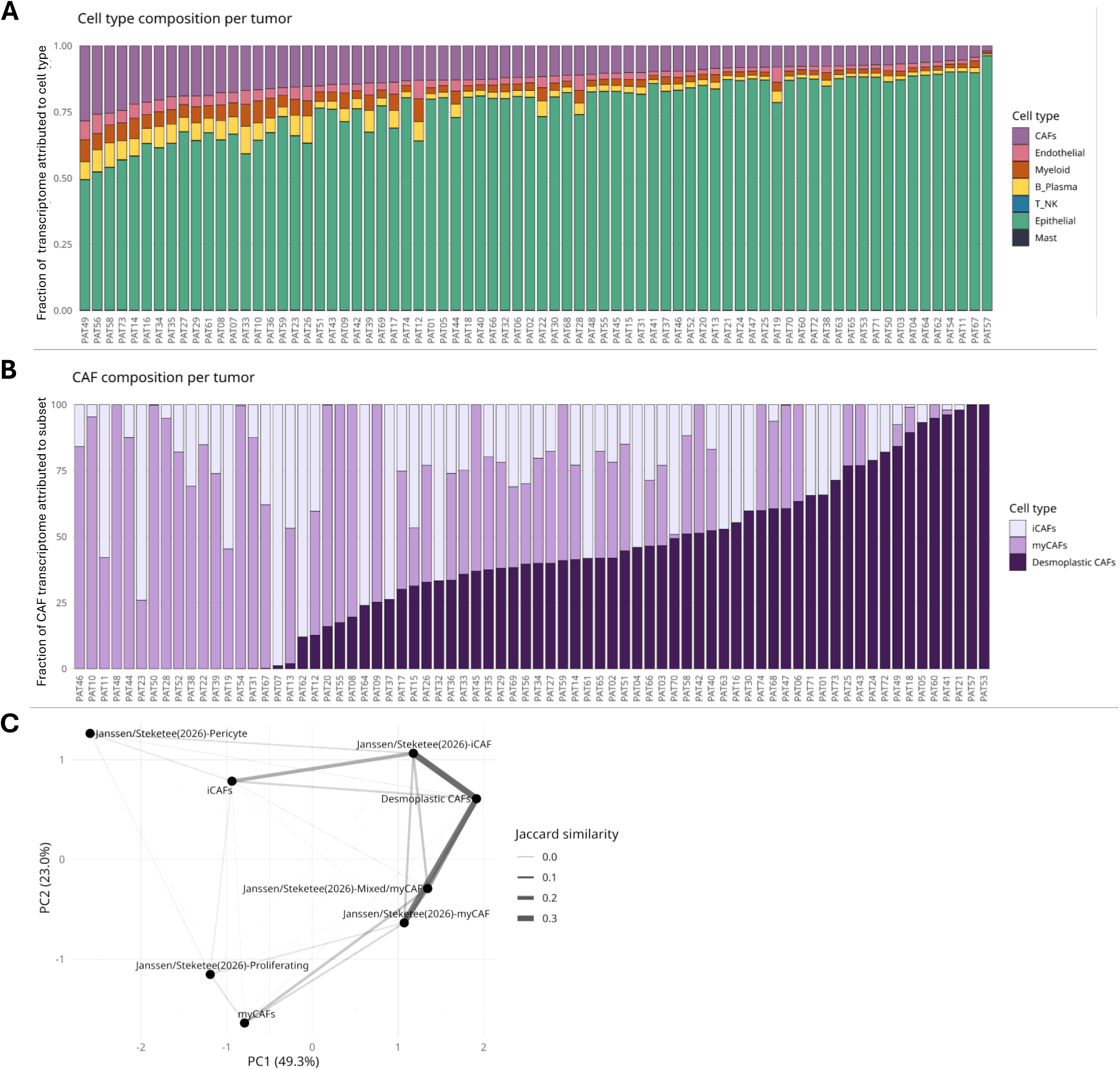
Identification of cell types in the PDAC tumor microenvironment using Statescope deconvolution. **A** Identification of different cell types using Statescope deconvolution in n=74 PDAC patients. PAT= pancreatic tumor **B** Identification of different CAF subsets using Statescope deconvolution, as a ratio to the total CAF population. PAT= pancreatic tumor **C** Principal component analysis comparing previously identified CAF subsets to the subsets identified in this cohort. Jaccard similarity index was displayed to show similarities between CAF subsets.

Each major cell type was further split into transcriptionally distinct cell states. Cell states were annotated based on the analysis of genes driving them. Epithelial cells were clustered based on classical or basal gene signatures as previously published [17] and further annotated using gene set enrichment (**Supplementary figure 1B-E, Supplementary figure 2, Supplementary table 2, Supplementary table 3**). Since the classical gene signature is characterized by high expression of epithelial genes, and the basal subtype is known for a high stromal content [18], we examined whether classical/basal subtyping correlated with the fraction of tumor cells present. However, the epithelial content was not higher among patients with more classical subtype epithelial cells (**Supplementary figure 3A**). This may reflect bias in the selection of tumor-rich regions used for NGS, or it may be because epithelial tumor cells undergoing epithelial-to-mesenchymal transition (EMT), a characteristic of the basal subtype, are still classified as epithelial cells in the deconvolution model. As expected, patients with a higher ratio of classical to basal epithelial cells had better disease-free and overall survival, supporting the reliability and validity of our data (**Supplementary figure 3B**).

Three CAF subsets were identified, which were classified as inflammatory CAFs (iCAFs: positive associating genes *PDGFRA*, *IL6ST*, *COL14A1* and *CXCL12*, and negative associating genes *ACTA2*, *TAGLN* and *COL1A1/2*), myofibroblastic CAFs (myCAFs: positive associating genes *MMP13/14*, *ADAM12* and *ACTG2*, negative associating gene *PDGFRA*) and desmoplastic CAFs (positive associating genes *COL1A1/2*, *SPARC* and *FBN1*) (**Figure 1B, Supplementary table 3**). Interestingly, these CAF subtypes showed some degree of similarity to the five CAF subtypes previously identified and published using a different cohort (**Figure 1C**) [13, 14]. Surprisingly, the desmoplastic CAFs showed a high degree of similarity to both the iCAFs and myCAFs identified in the referenced study, whereas they were quite distinct from the iCAFs and myCAFs we identified. This indicates that the desmoplastic CAFs form a transcriptionally distinct group relative to the CAF subtypes identified in our study. In line with our expectations, the iCAFs and myCAFs identified in this study showed similarities to the previously identified iCAFs and myCAFs. Next, we further determined whether FOLFIRINOX influences CAF abundance and phenotype. However, comparing the total abundance of CAFs and their subsets in untreated and neoadjuvant-treated patients showed no significant differences(**Supplementary figure 4**).

### CAF subsets differentially associate with classical and basal epithelial subsets, while desmoplastic CAFs only contribute to the presence of different myeloid and B cell types

Correlating the abundance of CAF subsets to epithelial subsets showed that iCAFs positively correlated with the classical EMT low, KRAS high epithelial subset. On the other hand, a strong correlation of the myCAFs and desmoplastic CAFs with basal epithelial subsets was observed, specifically the basal, MYC signaling and basal EMT high population, respectively. In addition, desmoplastic CAFs were also associated with both mixed epithelial subsets (**Figure 2A**).

**Figure 2.**
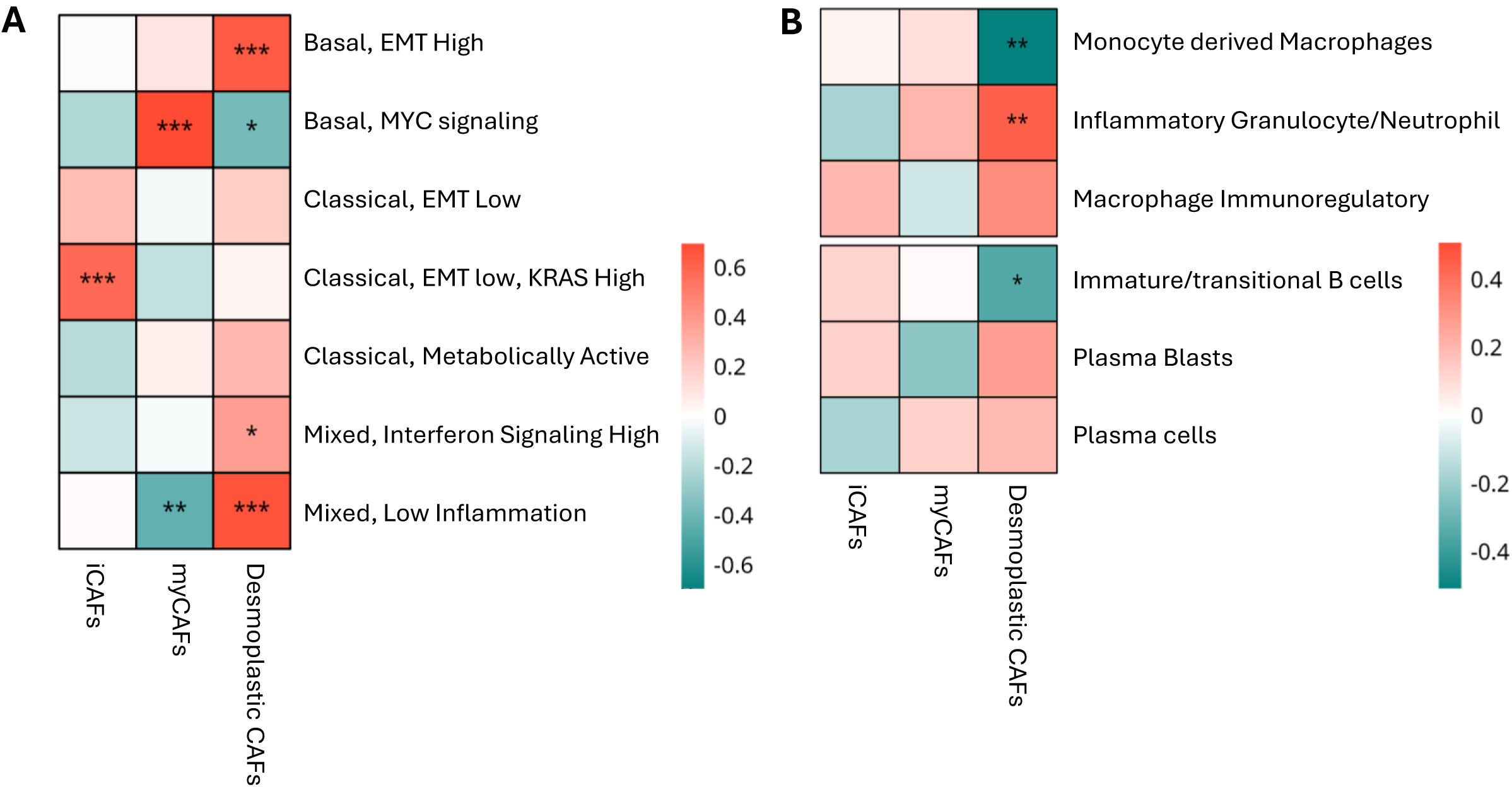
CAF subtypes correlate with the presence of different epithelial and immune cells. **A** Heatmap showing correlations between CAF subtypes and epithelial subtype abundance. **B** Heatmap showing correlations between CAF subtypes and myeloid cells/B and plasma cell abundance. Color gradient indicates strength and direction of correlation, stars indicate significance. *p<0,05, **p<0,01, ***p<0,001 (Benjamni Hochberg adjusted p values).

Myeloid cells constitute a substantial proportion of the immune cells in PDAC [21]. Although both iCAFs and myCAFs did not show any significant associations with the different myeloid subsets, desmoplastic CAFs showed a positive correlation with inflammatory granulocytes/neutrophils and a negative association with monocyte-derived macrophages (**Figure 2B**).

The same trend is apparent for the associations between CAF subsets and B and plasma cells, as both iCAFs and myCAFs also show no significant associations. On the other hand, desmoplastic CAFs showed a significant negative correlation with the immature and transitional B cells (**Figure 2B**).

### CAF subtypes do not associate with common PDAC mutations

Next, we analyzed the mutational spectra of the patients and correlated them with the identified CAF subsets. Mutations found in the cohort included common PDAC mutations, including KRAS (70/74 patients, 95%), TP53 (52/74 patients, 70%), CDKN2A (26/74 patients, 35%) and SMAD4 (10/74 patients, 14%). However, no correlations were identified between the mutational profiles and the abundance of CAF or epithelial subsets were identified (**Figure 3**).

**Figure 3.**
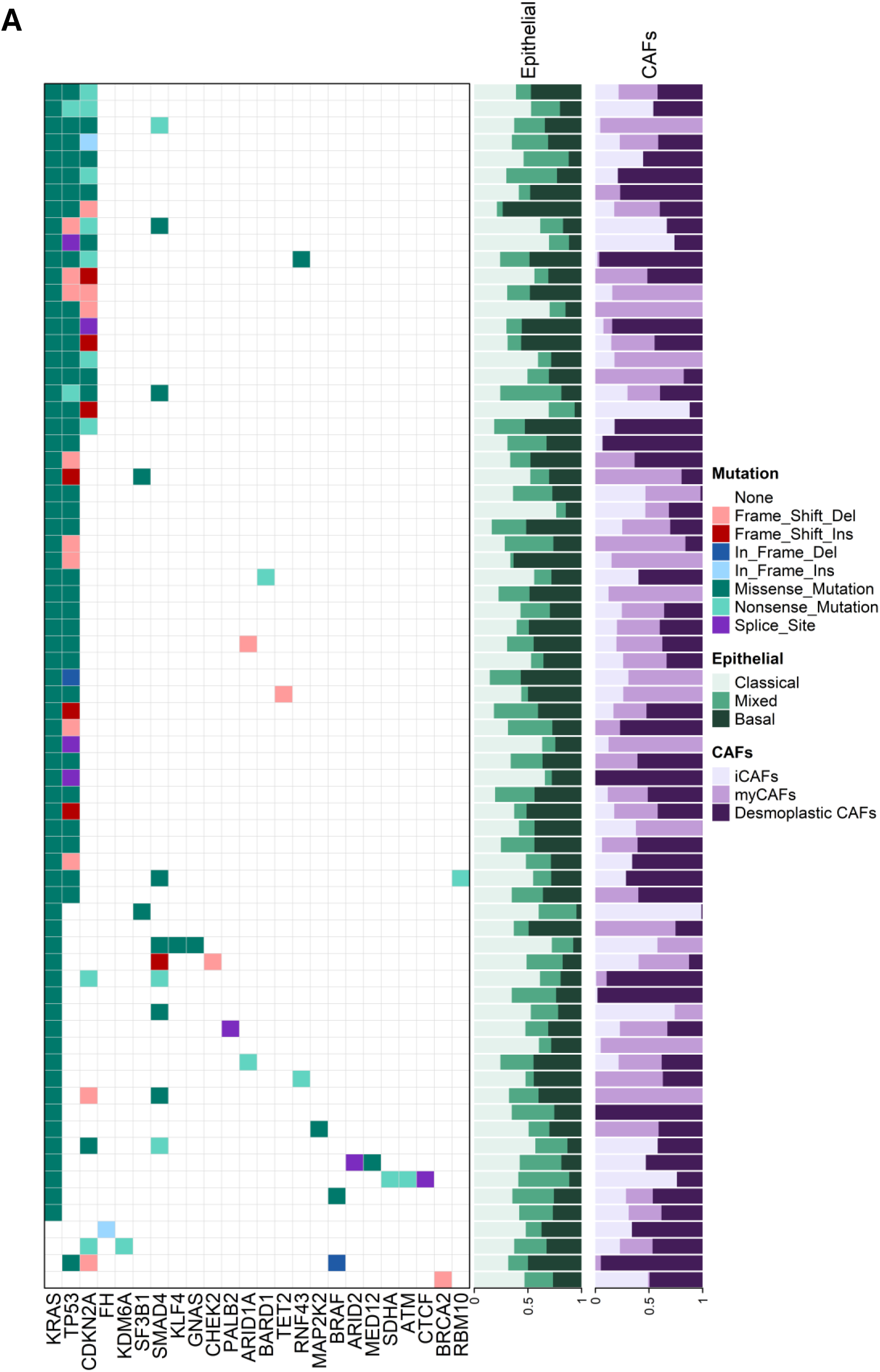
Overview of mutational signatures and CAF/epithelial subtype abundance. **A** Mutations occurring in each patient of the cohort, with epithelial and CAF subtype abundance displayed on the right. Different types of mutations are indicated with different colors.

### Increased abundance of total CAFs and iCAFs correlates to better survival in females

Lastly, the association of total CAF abundance and the different CAF subsets with disease-free survival and overall survival was determined, by splitting the cohort into two groups based on median abundance (compared to the total cells for total CAFs or to the total CAF population for CAF subsets). Surprisingly, a high abundance of total CAFs (CAF^high^) was associated with better overall and disease-free survival (OS and DFS, respectively, **Figure 4A**), while no significant differences in baseline characteristics were found between CAF^high^ and CAF^low^ patients (**Supplementary table 4**). This result is confirmed through univariate Cox regression modeling (OS: HR 0.4347 per 0.1 CAF increase, 95% CI 0.1980 to 0.8848, p=0.0205, DFS: HR 0.4832 per 0.1 CAF increase, 95% CI 0.2525 to 0.8774, p=0.0159). Interestingly, when total CAFs are stratified into CAF subsets, patients with higher iCAF abundance predominantly showed improved survival compared to those with low iCAF abundance, whereas myCAF or desmoplastic CAF abundance did not influence survival (**Figure 4B**). In univariate Cox regression modeling a similar trend was observed where iCAFs associate with improved survival, however this did not reach statistical significance (OS: HR 0.9305 per 0.1 CAF increase, 95% CI 0.7984 to 1.064, p=0.3065, DFS: HR: 0.9244 per 0.1 CAF increase, 95% CI: 0.8034 to 1.047, p=0.2248).

**Figure 4.**
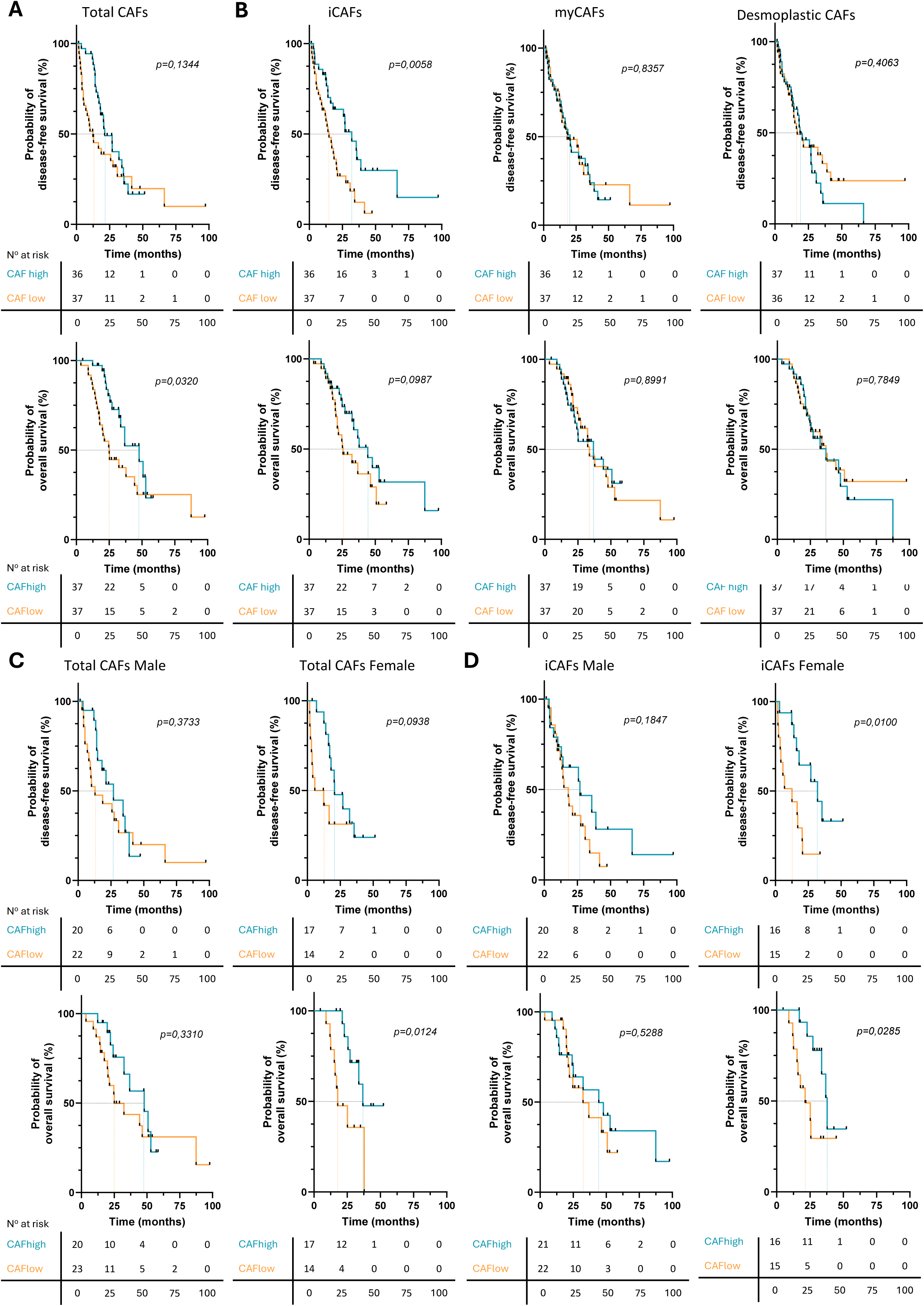
High total CAF and iCAF subtype abundance positively influence survival, especially in female patients. **A-D** Total CAF or CAF subtype abundance was split into high and low based on median abundance (CAF^high^ and CAF^low^, respectively), followed by Kaplan-Meier analysis of disease-free (top panel) and overall survival (lower panel). Patients were additionally stratified on gender in panels C and D. P-values were determined using the Log-rank (Mantel-Cox) test. The number of patients at risk at different time points is shown in the table below the Kaplan-Meier. Disease-free survival data were unknown for one patient. Dashed lines indicate median survival time. Events in overall survival includes death due to all causes, events in disease-free survival includes recurrence of the disease at any tumor stage (metastasis, locally advanced or local recurrence).

Recent literature has shown that female patients with PDAC have better survival than males, which has been associated with the presence of a favorable-prognosis iCAF subtype that is most commonly present in females [19, 20]. This iCAF subtype (termed iCAF.1) was characterized by high expression of the genes OGN and COL14A1, which are also in the top driving genes in our iCAF state (OGN loading iCAF: 0.09948, vs myCAF: -0.11971 and desmoplastic CAF: 0.05675, COL14A1 loading iCAF: 0.10795, vs myCAF: -0.14391 and desmoplastic CAF: 0.06620). In our cohort, females did not have better survival than males and no differences in CAF subtype abundance (or in most baseline characteristics) were observed between genders (**Supplementary figure 5A/B**, **Supplementary table 5**). However, analysis of survival data for total CAF and iCAF subtype abundance stratified by males and females showed that more iCAFs were prognostically relevant only in females, whereas no difference was observed in males (**Figure 4C/D**). This gender difference was not apparent for myCAFs and desmoplastic CAFs (**Supplementary figure 5C**).

## Conclusion and discussion

Until recently, single-cell RNA sequencing was considered the gold standard for identifying CAF subsets within a tumor. However, the use of this technique is limited by its cost and sequencing depth and introduces cell-type biases due to tissue dissociation and cell isolation protocols. Here, we instead used a deconvolution-based model called Statescope on bulk RNA sequencing data to identify cell subsets, including CAF subsets, within a large cohort of resected PDAC tumors. Deconvolution identified three CAF subsets, termed inflammatory CAFs (iCAFs), myofibroblastic CAFs (myCAFs) and desmoplastic CAFs. Our results have shown that the iCAF subtype is associated with the presence of classical epithelial cells, whereas myCAFs and desmoplastic CAFs correlate with different basal-subtype epithelial cells. The desmoplastic CAF associates with inflammatory granulocytes/neutrophils and negatively associates with monocyte-derived macrophages and immature B cells. Neither iCAFs nor myCAFs are associated with any of the identified immune cell clusters. In addition, no associations between the tumor mutational spectra and abundance of CAF/epithelial subsets were found. Interestingly, we found that higher CAF abundance was a strong prognostic indicator of improved survival, which seems primarily mediated through iCAF abundance and is predominantly apparent in female patients.

While the myCAF and iCAF subsets are well-characterized in PDAC, the desmoplastic CAF subset identified in this study is less well-known and has, to our knowledge, only been described in other tumor types [11, 12, 22]. In PDAC, myCAFs have been described as the CAF subtype responsible for desmoplasia, suggesting that in most PDAC datasets myCAFs and desmoplastic CAFs have been pooled into a single subtype. The reason that two subtypes are now identified could be influenced by spatial localization within the tumor, as myCAFs have been described to be closely located to tumor cells [23] and the samples used for this study were tumor-rich. In contrast, antigen-presenting CAFs (apCAFs), which have been previously studied in the context of PDAC [12], were not identified in this study. However, this population is sparse and markers that define this subset have previously clustered within the iCAF population of a human scRNAseq dataset, implying that apCAFs could also be clustered within the iCAF population in this dataset [12].

Previously, others have also classified CAF subsets in a larger cohort of PDAC tumors by applying a CAF subtype-specific gene signature to bulk RNA-seq data. This led to the identification of a so-called permissive and restraining CAF subtype (permCAF and restCAF, respectively) [24]. However, this model is limited in its ability to identify the abundance of distinct cellular subsets coexisting within a tumor and will classify the whole tumor as either permCAF or restCAF subtype. Statescope deconvolution allows detection of intratumoral heterogeneity within both the tumor cell compartment and the tumor microenvironment. In the cohort studied in this manuscript, high heterogeneity was observed across tumors from different patients, with most cell types varying substantially in abundance. This high inter-patient heterogeneity across cell types in PDAC has also been observed in previous studies [25–28]. The exception to this is the virtual absence of T and NK cells and mast cells in all samples, as observed in PDAC before [7, 29]. Like the high heterogeneity observed within cell types, heterogeneity in abundance is also observed for the different cell states. Using a different PDAC cohort (TCGA-PAAD), Statescope deconvolution previously identified five CAF subsets, whereas it identified three distinct CAF subsets in our cohort. Jaccard similarity indicates that our CAF subtypes show some similarities with the previously identified CAF subtypes. In addition to identifying a different number of CAF subsets, Statescope applied to the TCGA-PAAD cohort identified only four tumor cell clusters, including a normal epithelial compartment, whereas we identified seven tumor cell clusters and no normal epithelium. Similar to the previous study, we have identified tumor cell types that can be clustered into basal- or classical-like epithelial cells. The differences in the identification of cellular states, both in the CAF and epithelial compartments, could be attributed to intrinsic differences between the two cohorts. For instance, patients in our cohort are aged 60 years and younger (median age at diagnosis 55 years), whereas the median age of diagnosis in the TCGA-PAAD cohort was 65 years [30]. It has been shown that ageing can alter the phenotype of CAFs, specifically by increasing their secretion of growth factors [31]. This could contribute to the identification of the proliferating CAF subset in the TCGA cohort, which was not identified in our study. Furthermore, the percentage of tumor cells was lower in the TCGA-PAAD cohort than in our cohort [30], which could explain why we identified more tumor cell clusters than CAF clusters.

Analysis of the identified CAF clusters relative to other cell types in the TME shows that iCAFs are associated with good-prognosis classical-like tumor cells, characterized by an EMT-low and KRAS-high gene signature, whereas myCAFs and desmoplastic CAFs correlate with basal-like tumor cells. Basal PDAC tumors are characterized by desmoplasia and have often been associated with an abundance of myCAFs, consistent with the findings of this study. Similarly, iCAFs previously identified by scRNAseq studies, including the good-prognosis iCAF.1 subtype mentioned in the results, have been associated with classical-like tumor cells [20, 24, 35, 36]. Interestingly, the associations of the myCAFs and desmoplastic CAFs with the different types of basal and mixed epithelial subsets seem to be reversed, with the basal MYC signaling tumor subset positively associating with myCAFs, while negatively associating with desmoplastic CAFs. Similarly, the mixed low-inflammation epithelial subset negatively associates with myCAFs, whereas it positively associates with desmoplastic CAFs. This indicates that myCAFs and desmoplastic CAFs can exhibit opposite features that influence the TME.

In this dataset, three myeloid subsets have been identified using Statescope, each with distinct roles on the overall activity of the immune system, including inflammatory and immunosuppressive cell types. Monocyte-derived macrophages are often considered pro-inflammatory, but can also contribute to immunosuppression and therapy resistance in cancer [37]. In our dataset, this subset shows a predominantly inflammatory gene signature. Similarly, the gene signature that defines inflammatory granulocytes and neutrophils also appears to indicate immune activation. Interestingly, the desmoplastic CAFs show an opposite association with both inflammatory myeloid subsets. However, classifying these myeloid subsets as strictly inflammatory or immunosuppressive may not fully capture their role within the TME, as each subset likely comprises a heterogeneous mix of cell types. For instance, neutrophils are generally associated with a poor prognosis in PDAC, but can also be highly plastic and functionally diverse [38]. Therefore, spatial localization of the different markers and further subdivision is needed to further verify the role of CAF subsets in the immune cell landscape of PDAC.

We further found that common PDAC mutations are also abundant in this cohort, with most patients harboring KRAS and TP53 mutations, and CDKN2A and SMAD4 mutations in part of the cohort. The high frequency of KRAS and TP53 mutations in our cohort is consistent with previously reported frequencies in the literature. Furthermore, TP53 mutations often occur subsequent to KRAS mutations, as is also apparent by the co-occurrence of KRAS and TP53 mutations in most patients in our study [40, 41]. These findings therefore indicate that the molecular profile of our cohort is representative of patients with PDAC. However, no correlations have been found between patients’ mutational spectra and the abundance of CAF subtypes and classical/basal epithelial cells. While others have associated tumor budding, a characteristic of EMT, with CDKN2A and SMAD4 mutations [42], as well as a CAF-based gene signature associated with low clinical risk to higher frequency of KRAS mutations [43], we have not found these associations. However, tumor budding is an indirect measure of EMT occurrence, and the latter study did not quantify the abundance of CAFs and their subsets, contrary to our analysis. Furthermore, a review of available data indicates that PDAC mutations cannot fully account for the cellular heterogeneity observed in PDAC [44]. This observation is also supported by our data.

Surprisingly, our results indicate a positive contribution of higher overall CAF abundance to survival, even though an increased number of CAFs has often been associated with poorer patient survival [32, 33]. In particular, our data point to iCAFs as a determinant of this effect, as patients with higher iCAF abundance exhibited improved survival, whereas increased presence of myCAFs or desmoplastic CAFs showed no significant impact on prognosis. Although univariate Cox regression of iCAF abundance versus survival did not reach statistical significance, this could be due to the limited number of events and the high number of censored patients in our cohort (due to limited follow-up time), thereby limiting the power of this analysis. iCAFs have often been associated with a worse prognosis, as they are known to mediate immunosuppression, contrasting our results [34, 35]. However, recent publications have identified two subsets of iCAFs in PDAC that are associated with a favorable prognosis, termed iCAF.1 and interferon-regulated CAFs [19, 36]. The finding that iCAFs correlate with better survival is consistent with the permCAF/restCAF classification mentioned earlier, in which restCAF resembles the iCAF subtype and is also associated with a better prognosis [24]. Interestingly, in our cohort, high iCAF abundance is prognostically relevant only in females, which could indicate that the iCAF.1 subset (defined by OGN and COL14A1 expression, as also found in our iCAF subset) is enriched in the females of our cohort, as has also been shown previously [20].

Overall, this study identified a CAF population comprising three CAF subtypes in an early onset PDAC cohort of patients under 60 years old, each with distinct associations with survival and other TME components, including a sex-specific survival benefit linked to iCAFs. These findings highlight both the utility of Statescope deconvolution in capturing intratumoral diversity and complexity, as well as the functional heterogeneity of CAFs in early onset PDAC.

## Acknowledgements

The authors thank all patients for their contribution, and Merus NV for performing sequencing of this data.

## Author contributions

**Nicole Dam:** Conceptualization, Formal analysis, Investigation, Project administration, Visualization, Writing – original draft, Writing – review and editing. **Mischa F.B. Steketee:** Methodology, Software, Writing – review and editing. **Gaby S. Strijk:** Data curation, Resources, Writing – review and editing. **Willem de Koning:** Resources, Writing – review and editing. **Lukas J.A.C. Hawinkels:** Writing – review and editing. **Vera Kemp:** Funding acquisition, Writing – review and editing. **Casper H.J. van Eijck:** Conceptualization, Funding acquisition, Writing – review and editing. **Yongsoo Kim:** Methodology, Software, Writing – review and editing. **Casper W.F. van Eijck:** Data curation, Funding acquisition, Resources, Writing – review and editing. **Bram W. van Os:** Formal analysis, Investigation, Supervision, Visualization, Writing – review and editing.

