## Supplementary figures and tables for "Bulk RNA sequencing deconvolution of pancreatic ductal adenocarcinoma identifies cancer-associated fibroblast subsets associated with survival and tumor microenvironment composition"

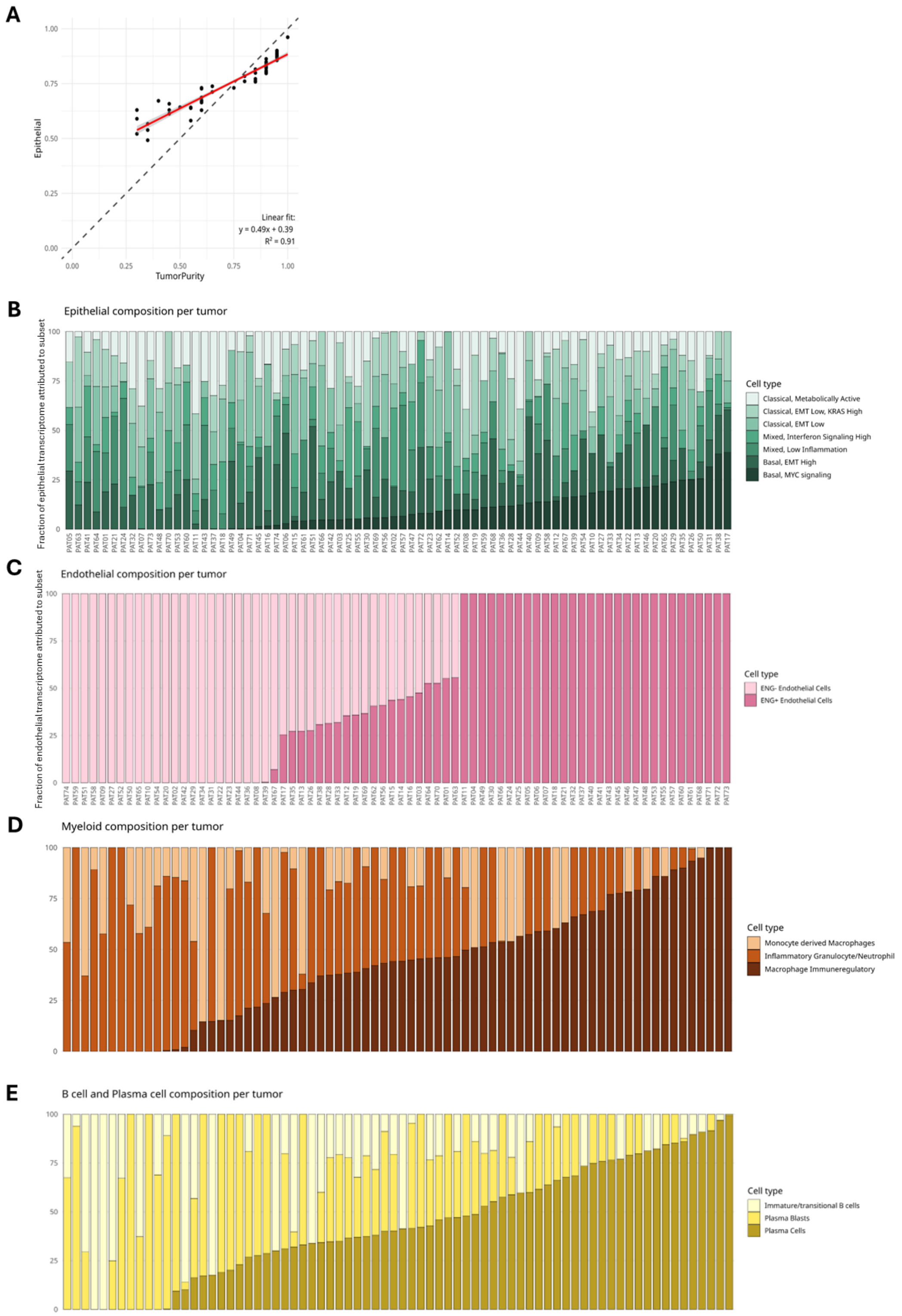


**Supplementary figure 1.** Identification of cell states. **A** Comparison of the epithelial cell fraction identified through Statescope (y-axis) to the tumor purity as identified through copy number variation (CNVkit). **B-E** Identification of different epithelial/tumor (B), endothelial (C), myeloid (D) and plasma/B subsets (E) using Statescope, as a ratio to the total population of each cell type. Cell subsets were named through analysis of the genes driving each subset, and for epithelial cells additionally through gene set enrichment as shown in supplementary figure 2. PAT= pancreatic tumor

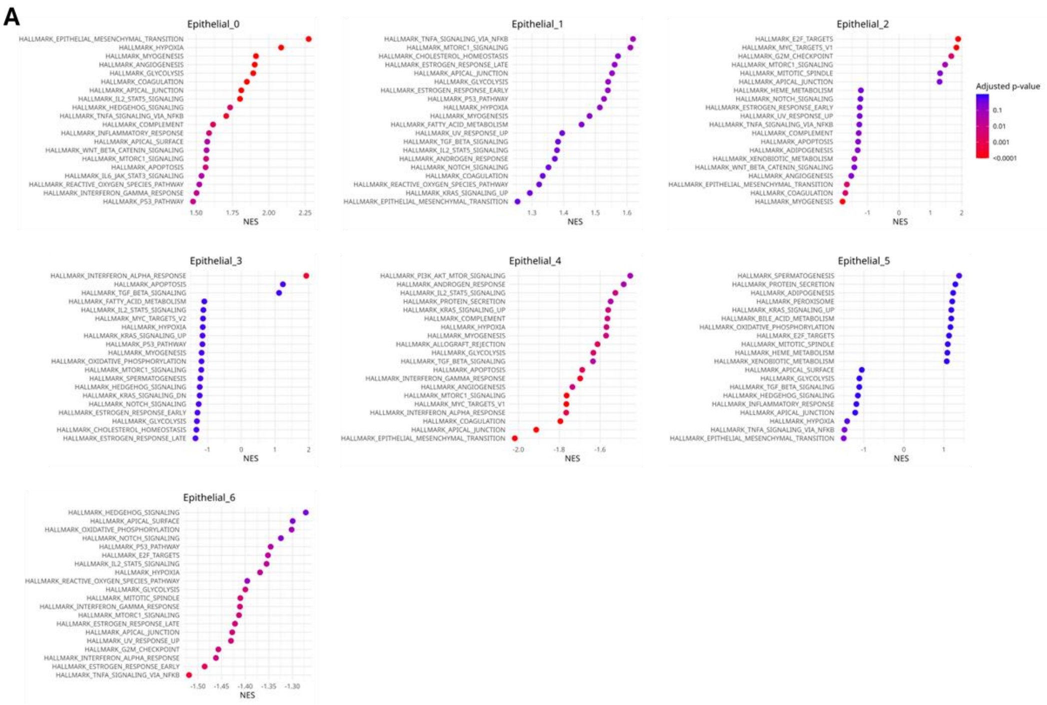


**Supplementary figure 2.** Identification of epithelial/tumor cell states **A** Gene set enrichment of epithelial cell states as shown in supplementary figure 1B.

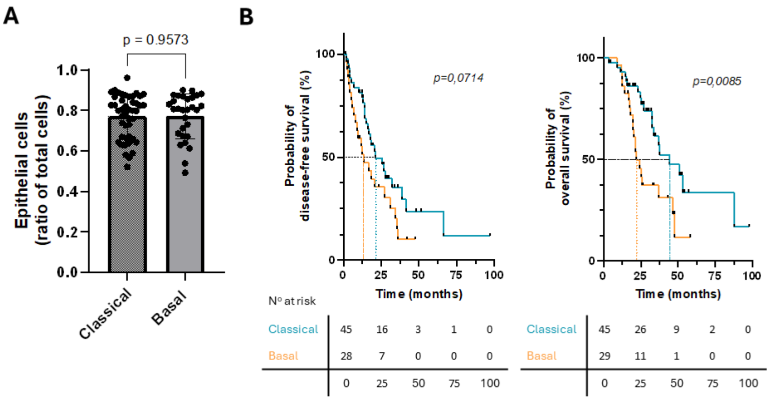


**Supplementary figure 3.** Epithelial cell states do not influence epithelial content but do influence survival. **A** Ratio of epithelial cells as a fraction of total cells against classical/basal tumor classification. Classical/basal classification was based on the most abundant subtype within the tumor. Significance was calculated using an unpaired T-test with Welch's correction. **B** Kaplan-Meier curve analysis of disease-free (left panel) and overall (right panel) survival in classical versus basal tumors. P-values were determined through the Log-rank (Mantel-Cox) test. The number of patients at risk at different time points is shown in the table below the Kaplan-Meier. Disease-free survival data were unknown for one patient. Dashed lines indicate median survival time.

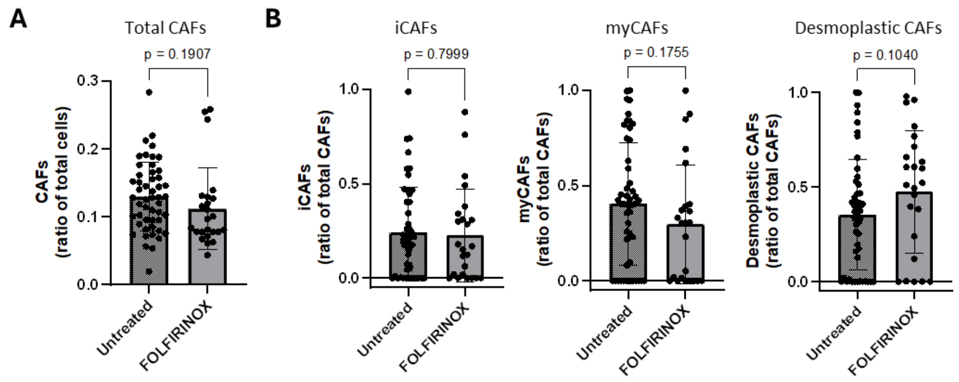


**Supplementary figure 4.** Total CAF and CAF subtype abundance are not influenced by neoadjuvant FOLFIRINOX treatment. **A** CAF abundance as a ratio of total cells compared between untreated and neoadjuvant FOLFIRINOX-treated patients. Significance was calculated using an unpaired T-test. **B** CAF subtype abundance (iCAFs, myCAFs, desmoplastic CAFs) as a ratio of total CAFs compared between untreated and neoadjuvant FOLFIRINOX-treated patients. Significance was calculated using an unpaired T-test.

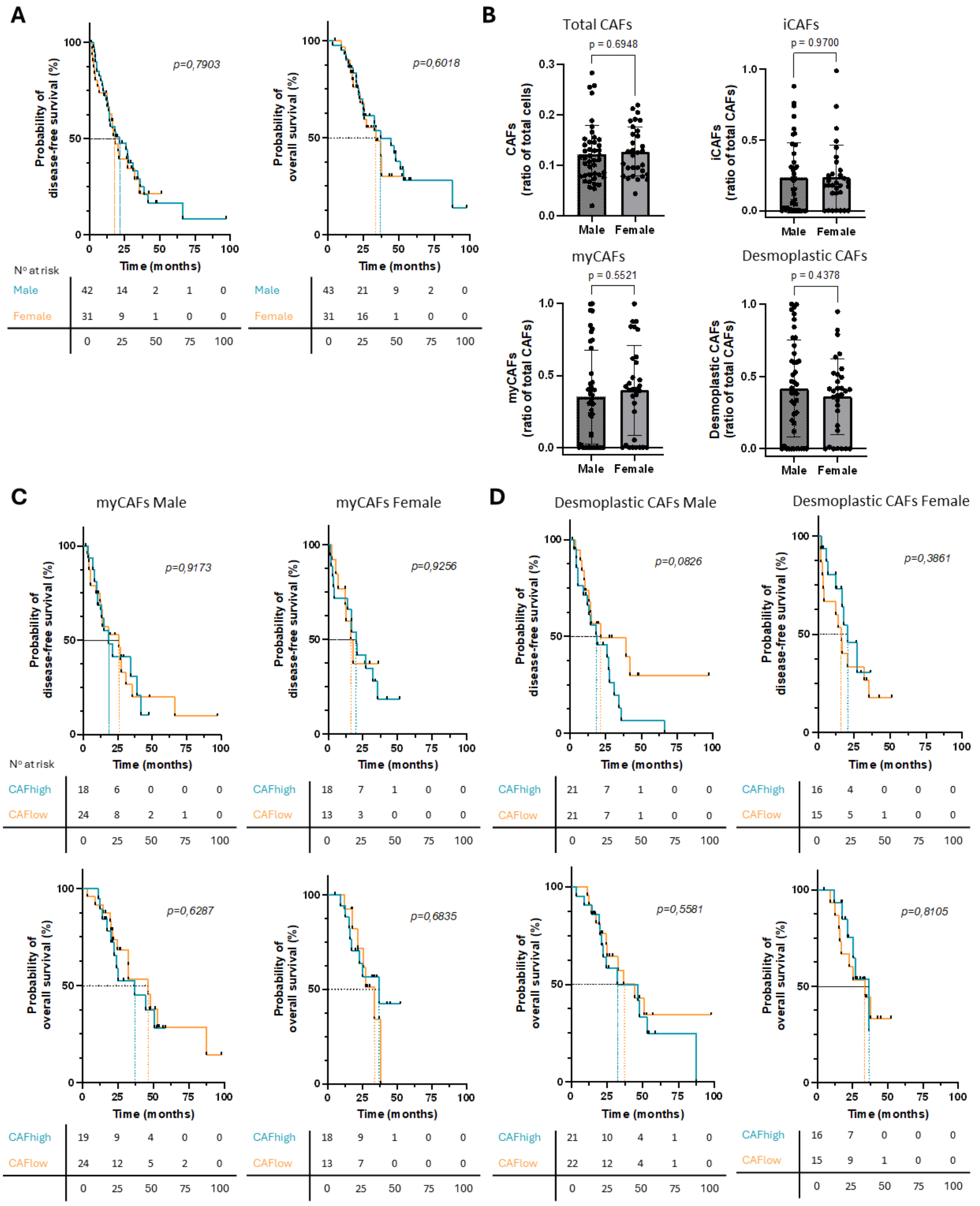


**Supplementary figure 5.** Males and females do not differ in their survival chances or CAF abundance. **A** Kaplan-Meier curve analysis of disease-free (left panel) and overall (right panel) survival in females versus males. P-values were determined through the Log-rank (Mantel-Cox) test. The number of patients at risk at different time points is shown in the table below the Kaplan-Meier. Disease-free survival data were unknown for one patient. Dashed lines indicate median survival time. **B** Total CAF, as ratio of total cells, and CAF subtype, as ratio of total CAFs, abundance in males and females. Significance was calculated using an unpaired T-test. **C/D** Kaplan-Meier curve analysis of disease-free (top panel) and overall (lower panel) survival based on myCAF (C) and desmoplastic CAF (D) abundance, split between males and females. P-values were determined through the Log-rank (Mantel-Cox) test. The number of patients at risk at different time points is shown in the table below the Kaplan-Meier. Disease-free survival data were unknown for one patient. Dashed lines indicate median survival time. Events in overall survival include death due to all causes, events in disease-free survival include recurrence of the disease at any tumor stage (metastasis, locally advanced or local recurrence).

### Supplementary tables

**Supplementary table 4.** Baseline characteristics of included patients, split by CAFhigh or CAFlow, n (%)

| | **CAFhigh (n=37)** | **CAFlow (n=37)** | p-value |
|---|---|---|---|
| Age diagnosis (median, IQR) | 55 (52-57) | 55 (49-57) | p=0,5534 |
| CA 19-9 at diagnosis (median, IQR) | 94 (24-240) | 70 (32,5-286) | p=0,9495 |
| Differentiation grade | | | |
| *Well* | 3 (8%) | 4 (11%) | p>0,9999 |
| *Moderate* | 14 (38%) | 15 (41%) | p>0,9999 |
| *Poor* | 11 (30%) | 9 (24%) | p=0,7940 |
| *Not assessed* | 9 (24%) | 9 (24%) | p>0,9999 |
| Neoadjuvant chemotherapy, FOLFIRINOX | 8 (22%) | 16 (43%) | p=0,0811 |
| Adjuvant chemotherapy | | | |
| *FOLFIRINOX* | 29 (78%) | 25 (68%) | p=0,4328 |
| *Gemcitabine* | 1 (3%) | 2 (5%) | p>0,9999 |
| *Gemcitabine + Capecitabine* | 0 (0%) | 1 (3%) | p>0,9999 |
| *None* | 7 (19%) | 9 (24%) | p=0,7784 |
| Disease stage at end of follow up | | | |
| *(borderline) resectable* | 0 (0%) | 0 (0%) | |
| *Locally advanced* | 0 (0%) | 0 (0%) | |
| *Metastatic* | 11 (30%) | 14 (38%) | p=0,6235 |
| *Resected* | 24 (65%) | 21 (57%) | p=0,6343 |
| *Local recurrence* | 2 (5%) | 2 (5%) | p>0,9999 |
| Death | 24 (65%) | 15 (41%) | p=0,0618 |

*IQR: interquartile range*

*NGS: next-generation sequencing*

**Supplementary table 5.** Baseline characteristics of included patients, split by gender, n (%)

| | **Male (n=43)** | **Female (n=31)** | p-value |
|---|---|---|---|
| Age diagnosis (median, IQR) | 54 (48,5-56) | 56 (52-58) | **p=0,0309** |
| CA 19-9 at diagnosis (median, IQR) | 106,5 (35,75-339,5) | 57,5 (21-144,75) | p=0,1124 |
| Differentiation grade | | | |
| *Well* | 3 (7%) | 4 (13%) | p=0,4430 |
| *Moderate* | 16 (37%) | 13 (42%) | p=0,8100 |
| *Poor* | 13 (30%) | 7 (23%) | p=0,5977 |
| *Not assessed* | 11 (26%) | 7 (23%) | p>0,9999 |
| Neoadjuvant chemotherapy, FOLFIRINOX | 16 (37%) | 8 (26%) | p=0,3273 |
| Adjuvant chemotherapy | | | |
| *FOLFIRINOX* | 29 (67%) | 25 (81%) | p=0,2900 |
| *Gemcitabine* | 3 (7%) | 0 (0%) | p=0,2597 |
| *Gemcitabine + Capecitabine* | 0 (0%) | 1 (3%) | p=0,4189 |
| *None* | 11 (26%) | 5 (16%) | p=0,3996 |
| Disease stage at end of follow-up | | | |
| *(borderline) resectable* | 0 (0%) | 0 (0%) | |
| *Locally advanced* | 0 (0%) | 0 (0%) | |
| *Metastatic* | 18 (42%) | 7 (23%) | p=0,1343 |
| *Resected* | 21 (49%) | 24 (77%) | **p=0,0164** |
| *Local recurrence* | 4 (9%) | 0 (0%) | p=0,1346 |
| Death | 24 (56%) | 15 (48%) | p=0,6383 |

*IQR: interquartile range*
*NGS: next-generation sequencing*
